# Building Genomic Analysis Pipelines in a Hackathon Setting with Bioinformatician Teams: DNA-seq, Epigenomics, Metagenomics and RNA-seq

**DOI:** 10.1101/018085

**Authors:** Ben Busby, Allissa Dillman, Claire L. Simpson, Ian Fingerman, Sijung Yun, David M. Kristensen, Lisa Federer, Naisha Shah, Matthew C. LaFave, Laura Jimenez-Barron, Manjusha Pande, Wen Luo, Brendan Miller, Cem Mayden, Dhruva Chandramohan, Kipper Fletez-Brant, Paul W. Bible, Sergej Nowoshilow, Alfred Chan, Eric JC Galvez, Jeremy Chignell, Joseph N. Paulson, Manoj Kandpal, Suhyeon Yoon, Esther Asaki, Abhinav Nellore, Adam Stine, Robert Sanders, Jesse Becker, Matt Lesko, Mordechai Abzug, Eugene Yaschenko

**Affiliations:** National Center for Biotechnology Information, National Library of Medicine, National Institutes of Health, Bethesda, Maryland, United States of America; Surgery, Center for Prostate Disease Research, Uniformed Services University of the Health Sciences, Bethesda, Maryland, United States of America; Computational and Statistical Genomics Branch, National Human Genome Research Institute, National Institutes of Health, Baltimore, Maryland, United States of America; Laboratory of Cell Biology, National Institute of Diabetes and Digestive and Kidney Diseases, National Institutes of Health, Bethesda, Maryland, United States of America; NIH Library, Division of Library Services, Office of Research Services, National Institutes of Health, Bethesda, Maryland, United States of America; Systems Genomics and Bioinformatics Unit, National Institute of Allergy and Infectious Diseases, National Institutes of Health, Bethesda, Maryland, United States of America; Translational and Functional Genomics Branch, National Human Genome Research Institute, National Institutes of Health, Bethesda, Maryland, United States of America; Stanley Institute for Cognitive Genomics, Cold Spring Harbor Laboratory, Cold Spring Harbor, New York, United States of America; Centro de Ciencias Genomicas, Universidad Nacional Autonoma de Mexico, Cuernavaca, Morelos, Mexico; Bioinformatics Core, University of Michigan, Ann Arbor, Michigan; Cancer Genomics Research Laboratory, Division of Cancer Epidemiology and Genetics, National Cancer Institute, National Institutes of Health, Gaithersburg, Maryland, United States of America; Department of Biology, Johns Hopkins University, Baltimore, Maryland, United States of America; Institute for Computational Biomedicine, Weill Cornell Medical College, New York, New York, United States of America; Tri-Institutional Training Program in Computational Biology and Medicine, New York, New York, United States of America; Department of Biostatistics, Johns Hopkins Bloomberg School of Public Health, Baltimore, Maryland, United States of America; McKusick-Nathans Institute of Genetic Medicine, Johns Hopkins University School of Medicine, Baltimore, Maryland, United States of America; Laboratory of Skin Biology, National Institute of Arthritis and Musculoskeletal and Skin Diseases, National Institutes of Health, Bethesda, Maryland, United States of America; Center for Regenerative Therapies, Technische Universität Dresden, Dresden, Germany; Translational Immunology, John Wayne Cancer Institute at Saint John’s Health Center, Santa Monica, California, United States of America; Microbial Immune Regulation Group, Helmholtz Centre for Infection Research, Braunschweig, Germany; Chemical and Biological Engineering, Colorado State University, Fort Collins, Colorado, United States of America; Graduate Program in Applied Mathematics & Statistics, and Scientific Computation, University of Maryland, College Park, Maryland, United States of America; Center for Bioinformatics and Computational Biology, University of Maryland, College Park, Maryland, United States of America; Division of Health and Biomedical Informatics, Department of Preventive Medicine, Feinberg School of Medicine, Northwestern University, Chicago, Illinois, United States of America; Genetics and Molecular Biology Branch, National Human Genome Research Institute, National Institutes of Health, Bethesda, Maryland, United States of America; Bioinformatics and Molecular Analysis Section, Center for Information Technology, National Institutes of Health, Bethesda, Maryland, United States of America; SRA International, Fairfax, Virginia, United States of America; Department of Computer Science, Johns Hopkins University, Baltimore, Maryland, United States of America

## Abstract

We assembled teams of genomics professionals to assess whether we could rapidly develop pipelines to answer biological questions commonly asked by biologists and others new to bioinformatics by facilitating analysis of high-throughput sequencing data. In January 2015, teams were assembled on the National Institutes of Health (NIH) campus to address questions in the DNA-seq, epigenomics, metagenomics and RNA-seq subfields of genomics. The only two rules for this hackathon were that either the data used were housed at the National Center for Biotechnology Information (NCBI) or would be submitted there by a participant in the next six months, and that all software going into the pipeline was open-source or open-use. Questions proposed by organizers, as well as suggested tools and approaches, were distributed to participants a few days before the event and were refined during the event. Pipelines were published on GitHub, a web service providing publicly available, free-usage tiers for collaborative software development (https://github.com/features/). The code was published at https://github.com/DCGenomics/ with separate repositories for each team, starting with hackathon_v001.

## Introduction

Genomic analysis leverages large datasets generated by sequencing technologies in order to gain better understanding of the genomes of humans and other species. Given its reliance on large datasets with complex interactions and its fairly regularized metadata, genomic analysis is an exemplar of “big data” science (1). Genomic analysis has shown great promise in finding actionable variants for rare diseases (2), as well as directing more specific clinical action for common diseases (3, 4).

Due to its potential for significant clinical and basic science discoveries, genomics has drawn many newcomers from the biological and computational sciences, as well as investigators from new graduate programs in bioinformatics. While many of these investigators can run established pipelines on local, public, or combined datasets, most do not have the expertise or resources to establish and validate novel pipelines. Additionally, highly experienced genomic investigators often lack the resources necessary to generate and distribute pipelines with broader applicability outside their specific area of research. In this study, we aimed to assess whether we could close this gap by bringing genomics experts from around the world together to establish public pipelines that can be both used by newcomers to genomics and refined by other seasoned professionals.

We sought to achieve this by hosting a hackathon at the National Institutes of Health (NIH) in Bethesda, Maryland. Hackathons are events in which individuals with expertise in various areas of software development and science come together to collaborate intensively over a period of several days, typically focusing on a specific goal or application. This form of crowdsourcing facilitates innovative ideas and agile development of solutions for challenging questions (5). Hackathon participants also benefit from the opportunity to learn new skills, network with other professionals, and gain the personal satisfaction of using their expertise to help the community.

Those of us who had previously attended hackathons had been motivated by learning from the experience, personal curiosity, professional networking, and the satisfaction of applying one’s skills to help the community. These previously attended hackathons included IT/programming-centric events, such as the Kaiser Permanente-sponsored hackathon for apps that could help prevent and fight obesity (6); bioinformatics-oriented events, such as the Illumina-sponsored hackathon to create apps in their proprietary BaseSpace cloud computing environment (7); and events focused on social issues, such as exploring women’s empowerment and nutrition in the developing world (8).

Hackathons are more common in software development communities than in bioinformatics and medicine, and may represent a valuable opportunity to accelerate biomedical discovery and innovation. Hackathons have gained popularity in the bioinformatics community in the last several years, though the environment and culture of a bioinformatics hackathon differs from that of typical IT/programming-centric hackathons (9). Bioinformatic hackathons more closely resemble a scientific discussion and provide an opportunity to learn and delve more deeply into specific areas. Typically a successful hackathon requires participants with both coding/programming skills and domain-specific knowledge (e.g. DNA-seq and RNA-seq).

Four teams of 4-6 participants came together for this hackathon to answer questions in the fields of DNA-seq, RNA-seq, metagenomics and epigenomics. The initial topics for exploration were based on questions bioinformaticians frequently ask computational cores. Massive amounts of data have been generated and made publicly available in these fields using high-throughput experimental technologies. Powerful computational pipelines are needed to handle such large datasets, and to help analyze the data to answer biomedical questions. In addition, these topics have seen a significant increase in publications in recent years, as demonstrated in Figure 1.

**Fig. 1:**
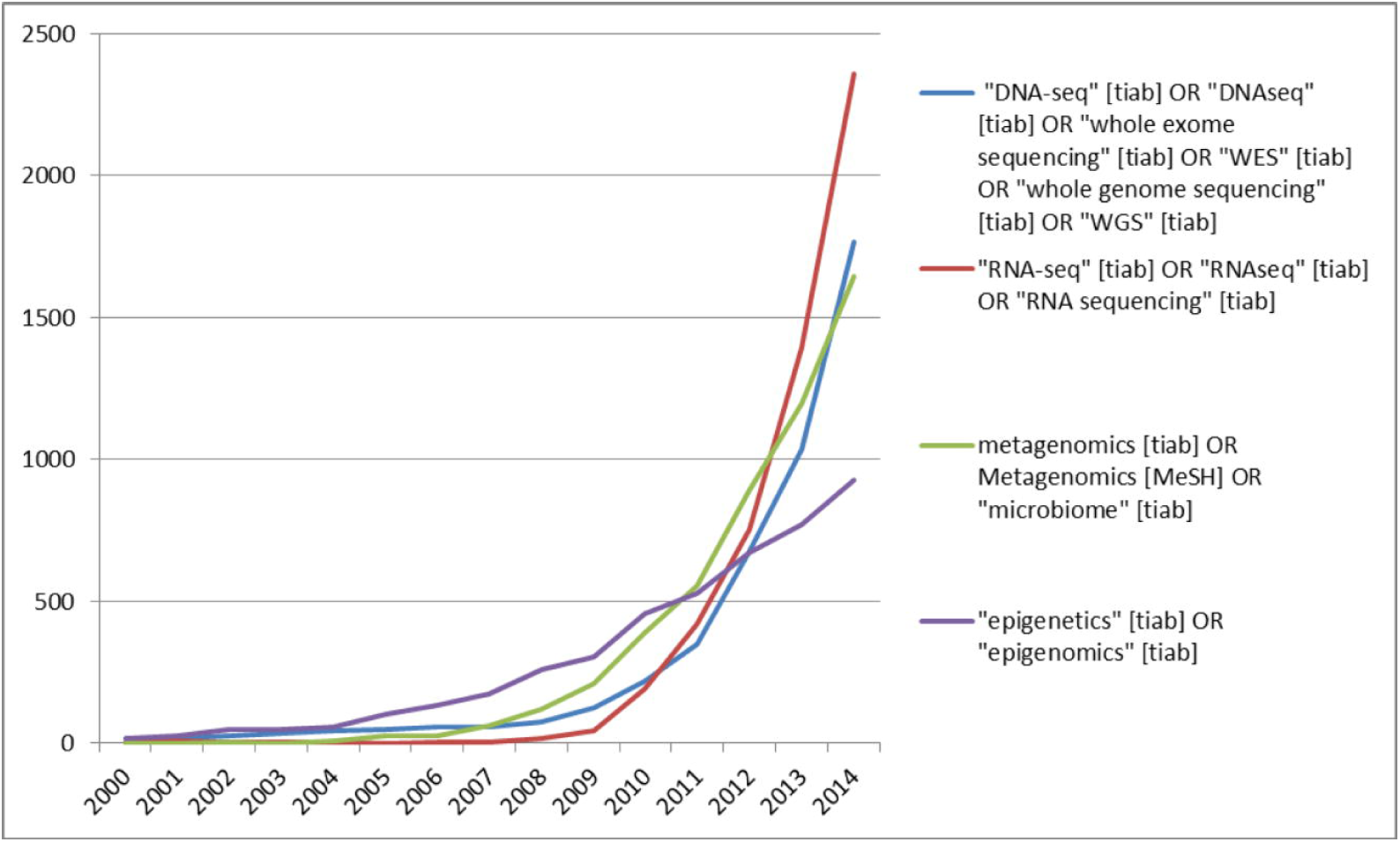
Articles indexed in MEDLINE for topics related to each of the four teams.

### Background for Hackathon Team Goals

#### DNA-seq Team

The goal for the DNA-seq Team was to create an easy-to-use integrated pipeline using existing tools to predict somatic variants from exome sequencing data, find shared and unique variants between samples, and filter and annotate the variants. Somatic mutation calling is particularly relevant to cancer genome characterization (10). Several methods exist to predict somatic variants by interrogating genomic sequences of tumor-germline paired samples (11–13). Issues with these variant calling algorithms are well known; each of the algorithms have strengths and weaknesses, and are sensitive to the datasets used (14). One way to test the reliability of the called variants and gain more confidence is to combine putative calls from several algorithms and use strict filtering criteria. Thus, we wanted to create a pipeline that included several existing calling algorithms. In addition, we intended to build a module within the pipeline to predict eQTLs using the called somatic variants and RNA-seq data from the same individuals. To achieve our goal of building an integrated pipeline, a paired exome sequence dataset as well as an RNA sequence dataset were required.

#### Epigenomics Team

Computational cores contain an abundance of epigenetic data encompassing a wide variety of markers over many different cell types. This abundance of information empowers labs to infer how these different markers contribute to gene expression and chromatin state. However, little standardization or collaboration exists among investigators seeking to elucidate relationships among epigenetic modifiers, leading to inconsistencies between analyses. The Epigenomics Team sought to analyze the extent to which DNA methylation and histone modifications affect gene expression using regression models. In addition to modelling, we wanted to create a framework for integrating experimental ChIP-seq or DNA methylation data provided by users into models of gene expression that were informed by publicly available data. A resource that allows users to make sense of their data in the context of the existing wealth of public data on epigenetic regulation provided an attractive and worthwhile goal.

#### Metagenomics Team

The Metagenomics Team worked on the problem of identifying the presence of viral sequences within human genomic or metagenomic sequences. The ability to locate these viral sequences offers the prospect of using computational tools for diagnostic purposes (15). Viral sequences may be embedded within human genomic sequence data as endogenous retroviruses or associated with bacterial consortia of the human microbiome as either lytic phages or prophages integrated into the bacterial genomes. Several studies have demonstrated the association between human-specific endogenous retroviruses (HERV) and diseases such as breast cancer (16, 17). In the context of the human microbiome, previous studies have used metagenomics to describe differences in bacterial consortia (18), but only a few studies have applied a metagenomics workflow to viral sequences present in human microbiome data (19, 20).

This team’s aim was to develop and compare analytical pipelines to identify and quantify viral sequences within human genomic and metagenomic datasets. The pipeline could be used in the characterization of the viral community in understanding the role viruses play in various environmental niches and diseases eventually associating differentially abundant viruses in disease or phenotype, despite database and sparsity issues potentially using marker-gene methodologies (21).

#### RNA-seq Team

RNA sequencing (RNA-seq) has become a useful technique for detecting gene expression levels, alternate splicing, and gene fusions (22). RNA variant detection can be important in cancer research to detect significant changes in tumor progression (23). The team’s initial goal was to build a variant calling pipeline using RNA-seq data from melanoma samples to differentiate germline and somatic mutations from RNA editing. Some sample datasets from the National Center for Biotechnology Information’s (NCBI) Sequence Read Archive (SRA) were suggested. Data could then be used to compare germline variants with known germline variants from the dbGaP version of the 1000 Genomes Project and ClinVar (24, 25). Another task was to determine isoform specificity based on mapping to 454 data using available software for the correlation with RNA structure prediction. Data could also be compared with the NCI-60 cell line variants. Also, we could try to determine mosaicism with germ-line vs intratumor data. Additional tasks were to detect systematic quality score variants indicating RNA editing with Illumina and Pacbio reads, correlate with RNA structure prediction and to fix HTseq annotation of tiny exons in UTRs.

## Materials and Methods

### Advertising, Preparation, and Logistics

Four team leads were selected by Busby to participate in the event. Team leads were experts in particular areas and they suggested scientific areas, datasets, or tools that they were familiar with. In order to recruit participants, we sent an announcement (Supporting Information to contacts in Busby’s network encouraging genomics professionals to apply for the hackathon, as well as posting it on several NCBI social media outlets, such as Facebook, Twitter, and Meetup.

After the application deadline, Busby reviewed the applicants for minimum necessary experience, and sent the applicants’ written statements to team leads, encouraging them to pick five members and two alternates. Team leads reviewed the 131 qualified applicants and selected a total of 27 people to fill the four teams of about 6 people each. Members were chosen solely on the interest and motivational statements on their forms, and credentials and experience were not further researched. One goal of the hackathon was to bring together scientists with different expertise. Some of the participants were more traditional bench biologists and some were bioinformaticists with more computational knowledge. This was a networking experience for local and international scientists, including two who traveled from Germany and several from outside of Maryland.

Technical professional staff from NCBI, including programmers and system administrators, were also brought in and “embedded” with the team to facilitate programming knowledge transfer and debugging. In addition, a librarian from the NIH Library served as an editor, providing guidance on writing and organization of the manuscript.

The teams convened in a large meeting space that had Wi-Fi access, a nearby cafeteria, plenty of tables to provide adequate room to spread out, and easels with markers for easier team discussion. Space was available outside of the meeting room if participants needed a quiet space apart from the larger group. To optimize time available to work on the projects, an hour was set aside each day for a “working lunch” with the option to order in as a group. There was also an optional group dinner each night that provided more time to network, socialize, and continue work on the scientific problems.

The schedule was circulated the week prior to the hackathon. The agenda was semi-structured with time allocated for various activities such as obtaining data, pipeline building, and code checking. There were several checkpoints for each team to present their progress to the larger group. A tour of the NCBI data center was conducted, as well as several brief informational presentations. For example, Eugene Yaschenko of NCBI presented a group of application program interfaces (APIs) relating to the SRA Toolkit collectively known as the software development kit (26). The schedule was designed to give team members a common starting point, but allowed for modification as needed for each team.

### Delegation of roles/responsibilities

One of the first tasks at the hackathon was for each team to assign roles/responsibilities to the team members. Most of the members were meeting each other for the first time, so it was a good way to learn about each other’s strengths. Roles were loosely distributed to cover areas including:

- Systems administration: decided where data and tools went, implemented git for versioning, and dealt with any technical issues;
- Metadata: tracked and recorded the information describing the contents of the datasets;
- Data Acquisition: located and downloaded appropriate publicly available datasets for analysis;
- Data Normalization: prepared data from multiple datasets to work in a pipeline, such as making read counts comparable between collection sites.
- Downstream Analysis: assigned annotation or function to genomic predictions and looked at statistically meaningful overrepresentation
- Documentation: prepared code and text summaries, including drafting this manuscript.

### Team Organization and Communication

In order to facilitate communication among team members, Google mail groups were created for each team. Most team members did not know each other prior to the start of the hackathon, but some teams used their Google mail group to communicate prior to the event. Other collaboration tools included GitHub for program version control and Gitter, an instant messaging tool, for sharing links, especially while teams were looking for datasets. Documents were collaboratively edited using Google Docs.

### Data Sources and Computational Tools

The basic workflow for the teams’ activities is demonstrated in Fig. 2. Scripting and data analysis took place in an Amazon Elastic Compute Cloud (EC2) where team players downloaded data from NCBI repositories and used open source bioinformatic tools. System administrators (SysAdmin) for each team were charged with code installation, compilation, storage and nodes expansion while the teams were developing the pipelines, which were pushed to GitHub.

**Figure 2.**
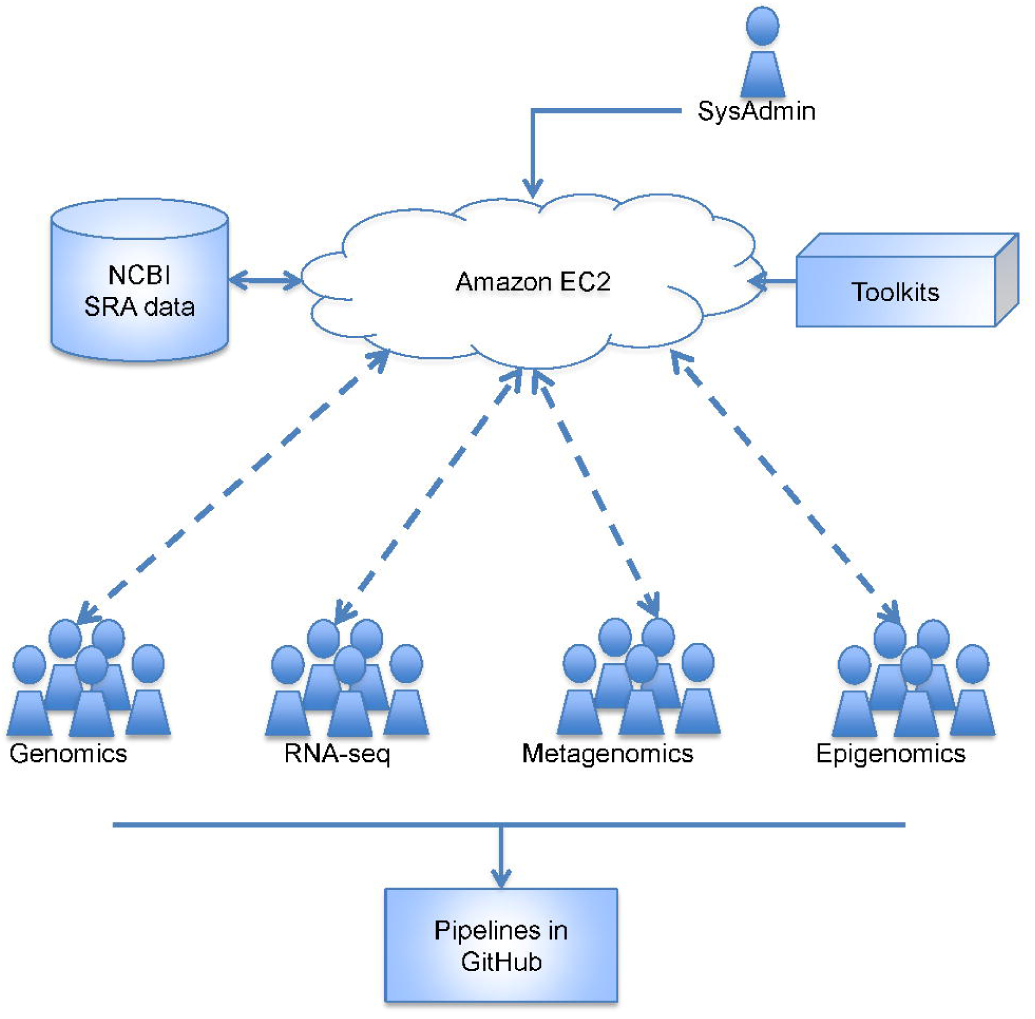
Hackathon teams workflow.

#### DNA-seq Team

The DNA-seq team acquired high-throughput sequencing data for matched tumor-normal pairs from the NCBI SRA. The first three datasets found were unusable due to corrupted files, mismatched samples, or missing header information (see Discussion below for more details). The data used for designing the DNA-seq pipeline were found by searching the BioProject database for the terms “homo sapiens NOT cell line,” filtering for exome and SRA datasets, and rejecting the 81 datasets that matched the term “HapMap”.

We designed our pipeline around a publicly available exome dataset submitted to SRA, BioProject PRJNA268172 (27). It consisted of exome sequences from four meningioma samples and a peripheral blood DNA sample from a 61-year-old female suffering from sporadic multiple meningiomas. Meningiomas are tumors originating from the membranous layers surrounding the central nervous system and are generally considered benign. Malignant meningiomas, while rare, are associated with a higher risk of local tumor recurrence and have a median survival time of less than two years (28). We were interested in finding and comparing somatic mutations, specifically SNPs, found in each of the meningioma tumor samples from the patient.

We downloaded the relevant files by using the SRA Toolkit, and converted the SRA files to SAM and BAM files for further analysis. We found that the SAM files contained reads with sequence and quality scores of different length, which would halt the BAM file conversion, so we added a filter in our pipeline to remove such reads. Ideally, a module for trimming/masking and re-aligning the FASTQ file would have been the most appropriate; however, due to time restrictions we were unable to add such a module to our pipeline. Here, we assumed that the SRA submitted files were properly aligned and followed appropriate quality control steps.

We used five different algorithms to call somatic variants. Four of these were in the Cake pipeline (29), namely Bambino (11), CaVEMan (30), SAMtools mpileup (31), and VarScan2 (12). To further increase our confidence in the called variants, we added the MuTect algorithm to those used by Cake. Xu et al have compared somatic variation calling algorithms (14).

A first level of filtering is provided in the pipeline for the resulted VCF files containing called somatic variants. This was accomplished using the Cake filtering module. We kept most of the default parameters, and those that were changed are explained in Table 1. Once the VCF files were filtered, they would be annotated using the ANNOVAR software, downloaded on January 5th, 2015 (32), with five different databases (RefGene, KnownGene, ClinVar, Cosmid and Cosmic), and the top 1% most deleterious CADD scores.

**Table 1:** Cake filtering module parameters modified by the DNA-seq Team.

| Parameter | Explanation | Flag or Value Used |
| --- | --- | --- |
| EXONIC_FILTER | Exome data is being used | FALSE |
| INDEL_FILTER | Only SNPs are being considered | FALSE |
| COSMIC_ANNOTATION_FLAG | Cosmic Annotations are added with ANNOVAR | FALSE |
| TUMOR_MIN_DEPTH | As the coverage of these samples was too low, we set the threshold to a less stringent value. | 5 |
| NORMAL_MIN_DEPTH | As the coverage of these samples was too low, we set the threshold to a less stringent value. | 5 |
| EMPTY_CONSEQ_FILTER | VEP annotations were not used | FALSE |

To compare the predicted somatic variations between two matched tumor-control samples, we created a module in our pipeline that used VCFtools to find shared and unique variations (33). Lastly, we combined these modules into a single pipeline using Bpipe (34), as well as a Unix Bash script. The full list of software used by the DNA-seq Team is included in Table 2.

**Table 2:** Tools and software used by the DNA-seq Team.

| Software/Tool | URL |
| --- | --- |
| SRA tool kit | http://www.ncbi.nlm.nih.gov/sra |
| Cake | http://sourceforge.net/projects/cakesomatic/ |
| MuTect | https://github.com/broadinstitute/mutect/releases/tag/1.1.5 |
| somatic sniper | http://gmt.genome.wustl.edu/packages/somatic-sniper/install.html |
| varscan | http://sourceforge.net/projects/varscan/ |
| bambino | https://cgwb.nci.nih.gov/goldenPath/bamview/documentation/index.html |
| picard | http://broadinstitute.github.io/picard/ |
| Annovar | http://www.openbioinformatics.org/cgi-bin/annovar_download.cgi |
| SAMtools | http://www.htslib.org/ |
| bpipe | http://github.com/ssadedin/bpipe |
| Python | https://www.python.org/ |
| GNU Parallel | http://ftp.gnu.org/gnu/parallel/ (35) |
| VCFTools | http://vcftools.sourceforge.net |

#### Epigenomics Team

We gathered data with the intention of modeling transcription (mRNA-seq) based on DNA methylation (RRBS or Bisulfite-seq) and histone states (ChIP-seq). To simplify analysis, we focused on marks associated with enhancers and their regulatory status: H3K27ac, H3K4me1, and H3K4me3. Ultimately, we required that included tissues have matching H3K27ac, RNA-seq and DNA methylation data for preliminary modeling. Data files were drawn from human cell lines and tissues in the NIH Epigenomic Roadmap that fit our criteria (outlined in Table 3), from the site’s FTP mirror (36) using the rsync command with the -av option. All files were already aligned, and were required to have been generated using the hg19 reference genome. Several files were identified that appeared to be re-aligned or uncorrected versions of other downloaded files, and were removed from the analysis. Tissues acceptable for analysis were identified by using the data matrix view on the Roadmap site, as well as searching for non-partial datasets with the Data Grid view of the International Human Epigenome Consortium (IHEC) Data Portal (37). Samples were initially downloaded from ENCODE: Encyclopedia of DNA Elements (38), but were not included in initial analysis for reasons of uniformity and time.

**Table 3:** NIH Epigenomic Roadmap Cell Lines/Tissues used by Epigenomics Team

| Cell/Tissue Type |  |
| --- | --- |
| H1 | Left_Ventricle |
| IMR90 | Pancreas |
| CD34_mobilized_primary_cells | Right_Atrium |
| hESC-derived_CD184+_endoderm_cultured_cells | Sigmoid_Colon |
| HUES64_cell_line | Spleen |
| pancreatic_islets | Penis_Foreskin_Fibroblast_Primary_Cells |
| Skeletal_muscle | Penis_Foreskin_Keratinocyte_Primary_Cells |
| Adrenal_Gland | Penis_Foreskin_Melanocyte_Primary_Cells |
| Aorta | Neurosphere_Cultured_Cells_Ganglionic_Eminence_Derived |
| Esophagus | Brain_Hippocampus_Middle |
| Gastric | Ovary |

#### Metagenomics Team

We designed six pipelines to compare different strategies for identifying and quantifying viral sequences among human genomic information. The pipelines are:

1. Identify and quantify HERV sequences in assembled reads using blastn,
2. Identify and quantify HERV sequences in non-assembled reads from a human genome using search tools from NCBI’s SRA Toolkit,
3. Identify and quantify HERV sequences in non-assembled reads from a human genome using a standard blastn search of reads in FASTA format.

Pipelines 4-6 repeat pipelines 1-3 to identify all viral sequences within a sample from a human bacterial microbiome.

For pipeline 1, whole genome sequence raw reads of human CEU NA12878 (39) were obtained from NCBI’s SRA database and converted from SRA to FASTQ file format using the fastq-dump command provided by the SRA Toolkit with default filter settings. The resulting FASTQ file was moved to an Amazon Elastic Compute Cloud (EC2) node and assembled into contigs with the ABySS assembler (40, 41). Contigs were used as queries for blastn against a database consisting of all *Retroviridae* RefSeq genomes that was constructed using the makeblastdb command. For pipeline 2, the SRA files containing raw reads for NA12878 were used directly as a database for a SRA blast (blastn_vdb) using the *Retroviridae* RefSeq genomes as query. For pipeline 3, the FASTA file of non-assembled reads from pipeline 1 was used as a query for a blastn search against the database of *Retroviridae* RefSeq genomes from pipeline 1.

For pipelines 4 – 6, we used samples from the Human Microbiome Project (HMP) that had passed preliminary quality checks (42). The plan for pipeline 4 was the same as for pipeline 1, except for using SOAPdenovo2 for the assembly of the microbiome FASTQ reads. For pipeline 5, rather than start with SRA files, raw Illumina WGS reads in FASTQ format first were converted to SRA format using the FASTQ loader tool latf-load within the SRA Toolkit. Pipeline 6 follows pipeline 3, but uses the microbiome FASTA as its query and the total viral RefSeq genome as its database. Full details about the data and tools we used are located in Tables 4 and 5, respectively.

**Table 4.** Data sources used by Metagenomics Team

| Data Source | FTP Location |
| --- | --- |
| refseq viral | ftp://ftp.ncbi.nlm.nih.gov/refseq/release/viral |
| 1000 genomes contigs | ftp://ftp.1000genomes.ebi.ac.uk/vol1/ftp/technical/reference/ |
| Human microbiome | ftp://public-ftp.hmpdacc.org/HMASM/PGAs/ |
| Human microbiome raw reads for right_retroauricular_crease | ftp://public-ftp.hmpdacc.org/Illumina/right_retroauricular_crease/SRS015381.tar.bz2 |
| Homo_sapiens_dna | ftp://ftp.ensembl.org/pub/current_fasta/homo_sapiens/dna/Homo_sapiens.GRCh38.dna.chromosome.10.fa.gz |

**Table 5:** Tools and software used by the Metagenomics Team.

| Software/Tool | URL |
| --- | --- |
| SRAToolkit | http://www.ncbi.nlm.nih.gov/Traces/sra/?view=software |
| ABYSS (from source, version 1.5.2) | http://www.bcgsc.ca/platform/bioinfo/software/abyss/releases/1.5.2 |
| R statistical package | http://www.r-project.org/ |
| BLAST+ | ftp://ftp.ncbi.nlm.nih.gov/blast/executables/blast+/LATEST/ |

#### RNA-seq Team

The RNA-seq Team determined that the samples initially suggested by the team lead were not appropriate, so we looked for paired tumor/normal datasets from the NCBI Sequence Read Archive (SRA). One dataset was a deep high-throughput transcriptome sequencing (RNA-seq) performed on three pairs of matched tumor and adjacent non-tumors (NT) tissues from HCC patients of Chinese origin, accession PRJNA149267 (43, 44) Another identified dataset was a study to identify a prognostic signature in colorectal cancer (CRC) patients with diverse progression and heterogeneity of CRCs, accession PRJNA218851 (45, 46). Thirty-six paired samples from this study (18 tumor samples: GSM1228184-GSM1228201 and 18 matched normal samples GSM1228202-GSM1228219) were also determined to suitable. Additional tools used by the RNA-seq Team are listed in Table 6.

**Table 6:** Tools and software used by the RNA-seq Team.

| Software/Tool | URL |
| --- | --- |
| HISAT | http://www.ccb.jhu.edu/software/hisat/index.shtml |
| Illumina iGenome | http://support.illumina.com/sequencing/sequencing_software/igenome.html |
| Bowtie | http://bowtie-bio.sourceforge.net/index.shtml |
| tophat | http://ccb.jhu.edu/software/tophat/index.shtml |
| Bambino | https://cgwb.nci.nih.gov/goldenPath/bamview/documentation/index.html |
| SAMtools | http://SAMtools.sourceforge.net/ |
| SRA Toolkit | http://www.ncbi.nlm.nih.gov/Traces/sra/?view=software |
| R | http://www.r-project.org/ |
| python | https://www.python.org/ |
| perl | http://www.perl.org/ |

## Results

### Evolution of the Projects

#### DNA-seq team

Our initial goal was to build a bioinformatics pipeline to predict somatic mutations using several calling algorithms and integrate the mutations with RNA-seq data to find eQTLs. However, one of the main hurdles we encountered was finding an appropriate dataset that could be used to test and help create our pipeline. Initially, we found a neuroblastoma dataset submitted to SRA that included both the exome and RNA-seq data for matched samples. However, the dataset was unusable because of corrupted files. The majority of our time at the hackathon was spent on finding an appropriate dataset and debugging issues with the publicly available files. This hindered our efforts to create a fully functional pipeline to achieve our goals. Due to a lack of a working dataset of both DNA-seq and RNA-seq from the same individuals, we were unable to write a module within our pipeline to find eQTLs. However, we were able to design a pipeline that would find somatic mutations using five calling algorithms for given matched samples using five different algorithms, filter and annotate mutations, and find shared and unique mutations between two matched sample pairs (see Methods for more details).

#### Epigenomics Team

We initially considered different scenarios in which a lab might utilize our proposed pipeline based upon their available datasets and how investigators might want to model their datasets. For example, investigators might want to find a relationship between DNA methylation levels and histone enrichment. Given the variation in epigenetic data available as part of publicly available datasets, we recognized the need for flexibility about which data components would be required to model different epigenetic relationships. Time limitations prevented us from generating a workflow for every potential scenario (for example, RNA-seq and ChIP-seq, or ChIP-seq and DNA methylation but no RNA-seq). Instead, we considered common questions that might interest a general epigenetics laboratory. Investigators with epigenetic data often want to understand how this data correlates to gene expression. Thus, we decided to focus our model on elucidating relationships between a given variable collection of epigenetic data and gene expression. A lab can use publicly available datasets to create a model with which to test their own epigenetic data.

#### Metagenomics Team

We considered several different options for metagenomics searches before the final six pipelines were settled. We spent significant time discussing the requirements for filtering or other QC steps on raw reads prior to running assembly (pipelines 1 and 4) or SRA blast steps. We also debated the relative merits of assembly for each task, eventually deciding to compare searches with and without assembly in different pipelines, which significantly added to our workload and may have contributed to the fact that only two pipelines were completed during the hackathon. With regard to assembly, our group considered using the established MG-RAST pipeline for a “brute force” blastn into raw reads (47), as well as using metAMOS (48), a comprehensive pipeline for metagenomics analysis for comparison. Eventually we decided to forego established pipelines, due to computational requirements within the timeframe of the hackathon and the desire to focus on developing novel workflows of our own.

For pipelines 1-3 we initially planned to use human genomic information from 1000 Genomes (24) but found the format (base genomic sequence with list of variants) more difficult to handle for our purposes than the human CEU NA12878 data that we eventually used. Likewise, for the microbiome task, we initially planned to apply our pipelines to several microbiome sample types, but eventually decided to focus on a skin microbiome, since skin samples tend to contain abundant viruses and multiple datasets may be available due to ease of sampling.

#### RNA-seq Team

Our original goals were very ambitious, especially the task of determining RNA editing without DNA controls. We were interested in looking at the variants in paired cancer samples, and spent a fair amount of time to find an appropriate dataset. We decided to align the samples using HISAT and determine the types and counts of variants (particularly A-I transitions) that may suggest RNA editing. We also wanted to determine variants in genes and then possibly correlate the genes to Gene Ontology.

We discussed a variety of aligners, such as STAR (49) and HISAT (50), which are very fast. Information about the recently released HISAT program was shared via the Google Group prior to the start of the hackathon so everyone had a chance to review this program. We selected HISAT because of its speed, a decision partly driven by the time constraints of the hackathon. We encountered some technical difficulties processing these in HISAT, so we ran the dataset with the 3 pairs of tumor/normal using tophat and bowtie so that we had some results for downstream processing, while another team member continued to develop the HISAT portion of the pipeline.

We selected bambino as our variant caller based on team members’ past successes with using this tool. A collection of Python and Perl scripts was written to filter out unmapped and low-quality reads and, more importantly, multi-mapped reads that did not map to a unique locus within the genome, since bambino would call each of these multiple alignments separately. The BAMs also had to be sorted by chromosome to prepare input for bambino. Bambino was then used to generate a variant call table and a Perl script was created to filter for coverage on both strands and give a sparser table for downstream analysis. Another program counted the variant info. We then applied the Fisher’s exact test to the data using R.

We discussed creating a command line to get gene ontologies of the set of genes, but were concerned about users keeping up-to-date versions of the GO database. Gene ontology could be determined by using web sites such as PANTHER (51) or DAVID (52, 53).

### Technical Problems

#### DNA-seq Team

We found the SRA website challenging to use for locating data, and the quality of available datasets was inconsistent. Although many data sources validate user-submitted files, a number of files that had been improperly validated and thus were not usable. Some files were corrupted and could not be used, such as those in BioProject PRJNA76777 (54). Our pipeline needed datasets that had a paired tumor-normal sample from the same patient. For some datasets, paired samples were not available, and other datasets were marked as being paired, when in fact they were not, such as BioProject PRJNA217947 (55). In addition, we observed that multiple datasets in the SRA database were missing the header information required to create BAM files used by downstream analysis tools, such as BioProject PRJDB1903 (56). In other cases, SRA data were found to be malformed, and caused certain tools to crash. Specifically, files from BioProject PRJNA268172 (27) contained reads with differing length sequence and quality scores (e.g. 34 bases of sequences, 70 bases of quality information). Files with such mismatches cannot be used in SAMtools to convert to BAM files, as a difference in these field lengths is inconsistent with the SAM format specification (57).

We also encountered problems with upstream bioinformatics code quality, such as poor or incorrect documentation. The tools we employed had a variety of installation methods, and few were available for easy installation through a package manager. For example, core software, such as R version 3, was not available as a package from the operating system vendor. Installing from a third-party repository is not complex, but may be daunting to someone inexperienced in systems administration.

#### Epigenomics Team

When searching for epigenetic datasets that belong to a given cell type, we found that in many cases all of the necessary data were not available in one centralized location. Thus, we had to search through multiple websites and databases to find enough epigenetic data for a given cell type we wanted to model. In some cases, the metadata for a given file was either corrupt or unavailable. In other cases, the assembly used to align reads for a given set of files was not clearly indicated, so these files were discarded. When dealing with wiggle (wig) and bigWig files, sometimes the format of the file was inconsistent and needed to be edited on the fly.

#### Metagenomics Team

Technical difficulties generally were resolved expediently, but still hindered timely analysis within the hackathon context. For example, some Amazon EC2 nodes would suddenly become completely unresponsive for unexplained reasons, requiring that we shut down and re-initiate the nodes. By the end of the hackathon, results were only available from the pipelines that used the SRA BLAST, in part because the SRA BLAST took about an order of magnitude less computing time than the standard blast program. In both cases, many Amazon compute nodes were available, but only the SRA BLAST was able to handle the large volume of human genome and human microbiome read data efficiently. In contrast, a huge amount of the processing power available to the standard blast program (several tens of nodes) was simply wasted while the program waited for data.

#### RNA-seq Team

It is important to recognize the difficulties of variant calling, especially with RNA-seq data. First, bias impacts genes expressed at lower levels. As gene expression itself varies from sample to sample, depth of coverage for any particular variant may differ. For instance, a variant in a sample with high gene expression would be called without difficulties, but may not be called in a sample that also carries the variant but whose expression is too low to call with confidence. Another source of variance lies within the heterogeneity of the tissue sample. Most tissue samples harbor multiple cell types, and not all of these cells will carry a somatic mutation. This problem is encountered in both DNA-seq and RNA-seq data, but results can be difficult to interpret on a per-variant basis when the fluctuation in overall coverage in gene expression is also considered. Thus, we decided to deal with overall global effects rather than selecting particular singular changes.

### Project Results

#### DNA-seq Team

The test dataset was downloaded from SRA website. The SRA Toolkit utility called prefetch allows the user to download SRA data files, but we found it initially troublesome to use due to configuration and storage issues; by default, prefetch stores all files in user home directories, which are often limited in storage capacity. We therefore wrote a faster web-scraper script to download the files from the SRA website. Given our time limitations, we had to rely on the user-submitted aligned and trimmed files, but we recommend that files submitted to the SRA should be validated prior to upload.

Our pipeline was designed to find somatic mutations using five different algorithms, filter and annotate the mutations, and compare the predicted mutations between matched tumor-normal samples. However, due to time constraints and initial difficulties with finding an appropriate data set and software installation, we were unable to complete our analysis. A diagram of the final DNA-seq Team pipeline design is presented in Fig 3.

**Figure 3.**
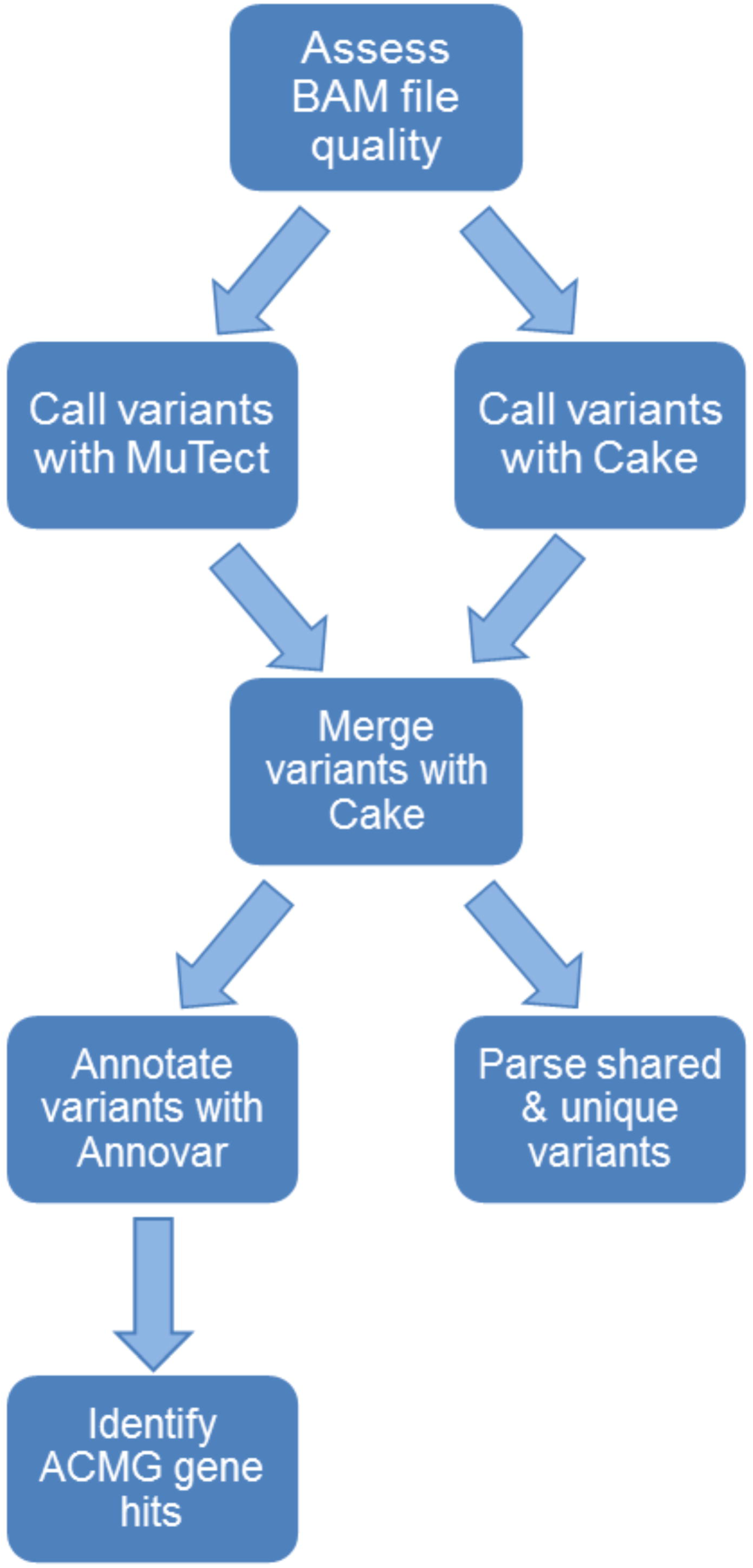
DNA-seq Team pipeline.

#### Epigenomics Team

We sought to rectify previously described inconsistencies in analyses by developing a more efficient, novel pipeline. Our pipeline uses RNA-seq counts, ChIP-seq peaks, and DNA methylation data in order to generate a model to predict relationships between gene expression and epigenetic data. These models can then be used to predict changes in gene expression with respect to changes in these epigenetic signals. Publicly available datasets can be utilized to generate a model, which investigators can then use to predict the state of the chromatin based on their own epigenetic data. The pipeline uses a combination of Python, R, and command line-based tools.

For each gene in a given cell type, epigenetic marks positioned locally to the gene are considered, as are distal enhancer elements that may also play in a role in that gene’s expression. To calculate the local epigenetic effects on transcription, an arbitrary distance on the 5’ and 3’ ends of a gene is binned into regions and the scores of epigenetic marks that reside in each of these bins are collected. The distal effect of transcription on a given gene is given by peak scores of enhancer elements that are at most one megabase (Mb) upstream or downstream of the gene.

The scores for each epigenetic mark and enhancer for a given gene are standardized and stored in a data matrix, where each row corresponds to a given gene for a given sample condition or cell type. Transcript gene counts generated from RNA-seq data are also stored. This pipeline generates a unique model for each gene in a given cell type by considering the gene count values as Y-values and each of the epigenetic scores as X-values. Corresponding coefficients are calculated for each X-value. Investigators can use these coefficients to input a new set of epigenetic data and receive a testable hypothesis of predicted levels of expression for each gene based on the new epigenetic data. Over time, different datasets can be used to train a given model to make it more reliable. A diagram of the final DNA-seq Team pipeline is presented in Fig 4.

**Figure 4.**
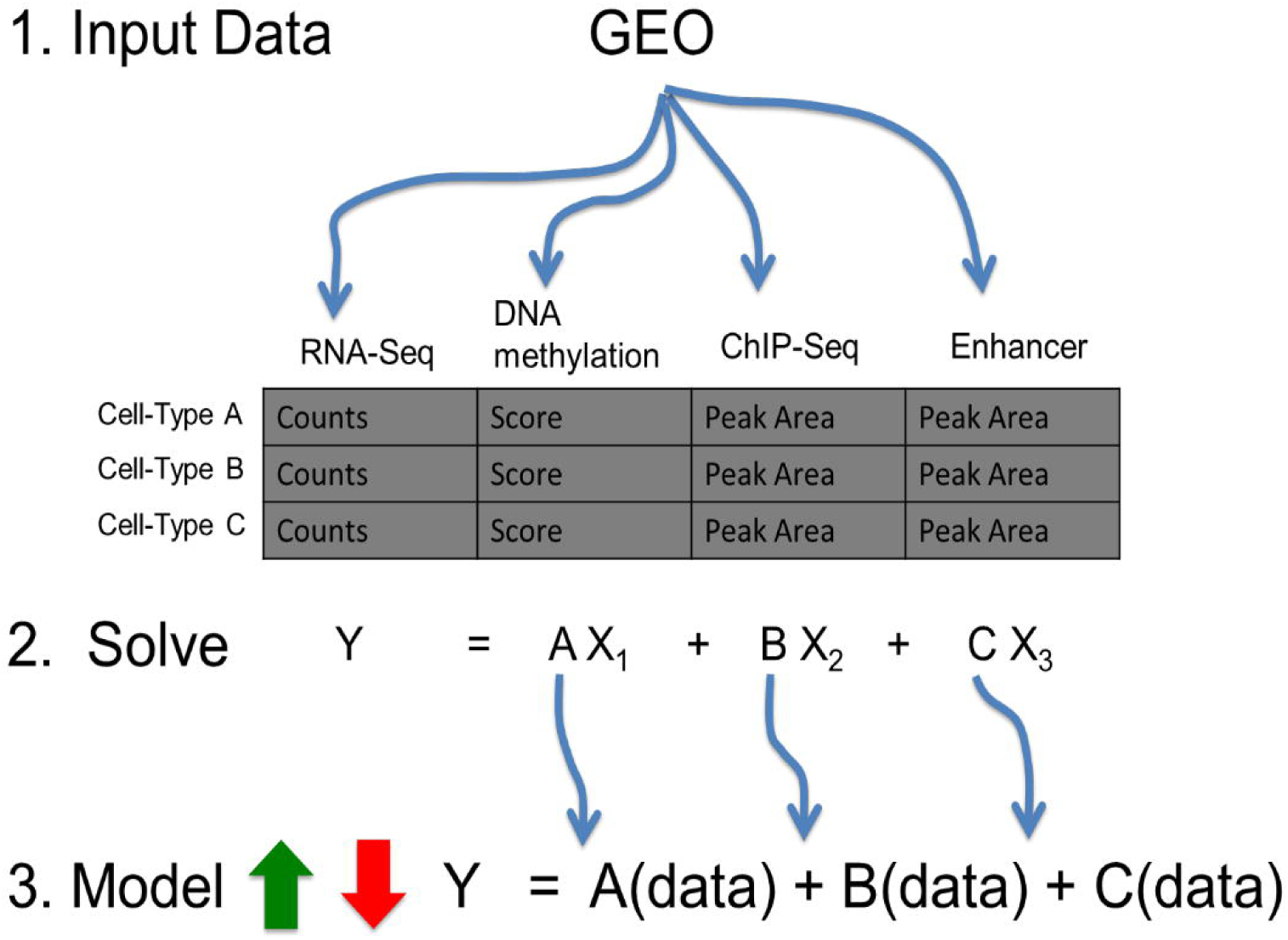
Epigenomics Team pipeline.

#### Metagenomics Team

Although the goals were similar across all six of our pipelines, differences in file formats and analysis approaches between the pipelines required the team to split their efforts rather than work together on a single pipeline. One result of this fragmentation was some lack of consistency in analytical methods (for example, choice of query versus database) between the pipelines. Moreover, due to time limitations of the hackathon only one assembly was completed: the ABySS assembly of the NA12878 human genome. Likewise, while we completed a versatile script for conversion of FASTQ files to SRA format with the latf-load tool, time allowed only for its demonstration on a single human microbiome sample.

Initially our plan included comparison of ERV sequence abundances between NA12878 genomes sequenced by several different sequencing technologies. Likewise, we initially planned to compare viral sequence abundances between several different microbiome sample types. Due to the complexity of these tasks, we decided to demonstrate our pipelines with a single sample type for each application: Illumina HiSeq 2000 reads from NA12878 and a single sample from the right retroauricular crease for the HMP application. Of the six pipelines that were planned and designed, we built four (pipelines 1-3 and 5).

We found that the most successful approach for searching a human genome for endogenous retroviruses was to use reads converted to SRA format (pipeline 2) via latf-load. The blastn in pipeline 2 was completed in 50-60 minutes. For pipeline 1, while an assembly of NA12878 was completed using ABySS within the time constraints of the hackathon, the blastn search using the assembled contigs to query the ERV database required excessive computational time; after more than 4 hours using 30 cores, the search still had not finished. In contrast, the blastn for pipeline 3 finished in 5 hours. Part of the increased time for the blastn search in pipeline 3 may have been due to alteration of the FASTA database by merging of forward and reverse paired-ends.

Pipeline 5 includes a set of scripts that we developed to create a versatile pipeline for searching a human microbiome sample for all viruses. These scripts may be adjusted to conduct BLAST searches using other types of SRA files. A shell script downloads the relevant datasets for the assembled and non-assembled sequences from HMP as well as for total viral sequences from RefSeq. Scripts and a wrapper, written in R, were developed to convert FASTQ data to SRA format with the latf-loader tool, convert the loaded data to .kar format, run a BLAST search with the blast_vdb command, and parse the data into a viral-by-sample count matrix. The resulting sparse matrix may be normalized and handled in a way similar to previously published methods for sparse matrices of high-throughput 16S survey data (21). Additional available code executes blastn on the assembled contigs. A diagram of the final Metagenomics Team pipeline is presented in Fig 5.

**Figure 5:**
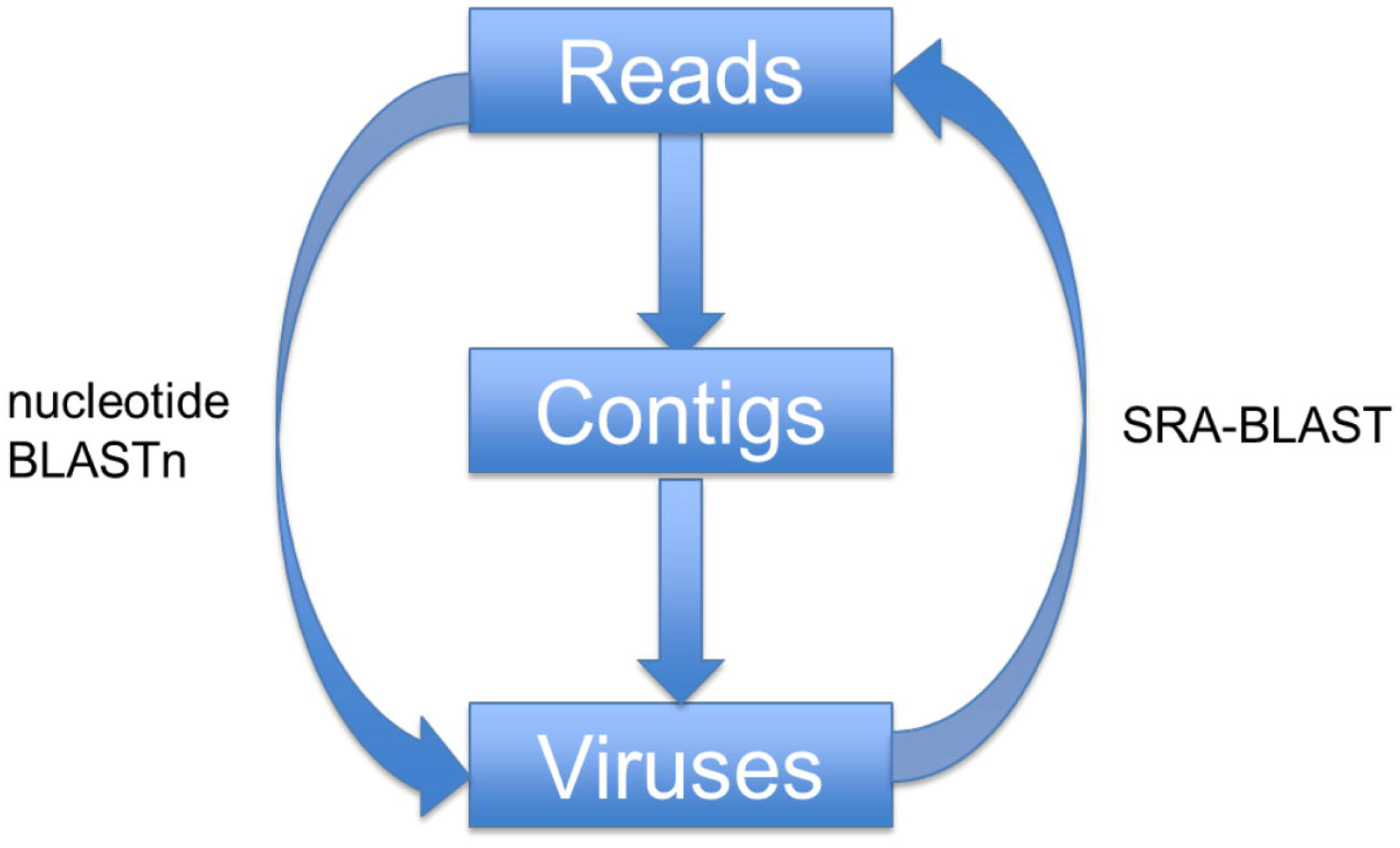
Metagenomics Team pipeline.

#### RNA-seq Team

We developed and ran a Python script that reads a user-defined manifest file to extract the read sequence information from the SRA files, stores the data in FASTQ format, and launches the jobs to align the sequences using HISAT. Due to technical difficulties and time constraints, we decided to manually download and process a smaller set of 3 pairs of tumor/normal samples, as opposed to the set of 18 pairs we had initially considered. We aligned the sequences using HISAT to prepare the data for use in subsequent parts of the pipeline. The aligned SAM files were filtered to remove the unmapped, low-quality or ambiguous reads, such as reads that map at multiple different locations.

The filtered data were run through bambino to create a variant call table in which each line contains a call variant at a particular location within the genome and the reference base at the same location. We counted nucleotide change variants in the tumor and normal samples and ran a Fisher’s exact statistical test using R to identify potential RNA editing. We found no significant global changes of overrepresentation, but it is important to recognize the limitations of our small sample size and our focus on specific changes. RNA editing most likely only comprises a small number of A to G variants, and we would not be able to identify these changes by considering global total numbers as opposed to looking at each site’s overall counts individually. This limitation does not affect overrepresentation in a global manner, but a small set of specific local changes might not be identified with this study design.

Before the end of the hackathon, we were able to use the initial Python script to download all 36 samples and launch the alignment tool jobs, but were able to complete fewer than 10 samples given the amount of time required to finish debugging. However, when completed, this automation script will greatly simplify the process of accessing and launching of alignment jobs for RNA-seq datasets. A diagram of the final RNA-seq Team pipeline is presented in Fig 6.

**Figure 6:**
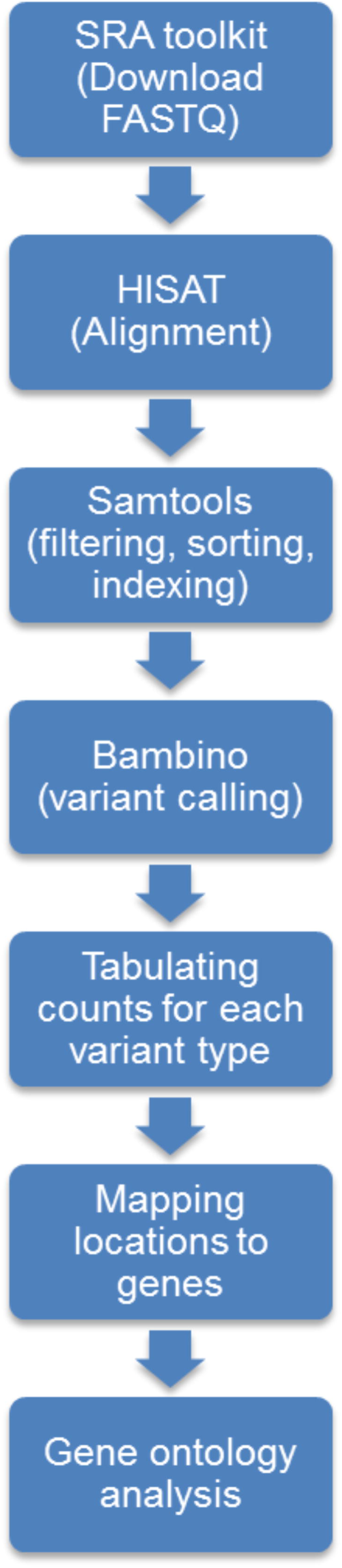
RNA-seq Team pipeline.

## Discussion and Conclusions

Feedback from hackathon participants was generally positive, and the enthusiasm that participants felt was evident during the event. Participants voluntarily stayed past the planned ending time each night, and many participants did not even want to take a break when lunch arrived. Even more than a week after the hacakathon had ended, teams continued to communicate about and work on the problems, as well as this paper.

Every participant in the hackathon contributed not only to the research but also to drafting the paper. Each group appointed a lead writer, who worked closely with the librarian editor and coordinated with the other members of their team. Because each of the team members worked on different parts of the project, every individual wrote at least a portion of the sections of the paper covering their work. The use of Google Docs allowed multiple authors to work on the paper simultaneously and all changes to be reflected in real time. Google Docs’ comment functionality also facilitated communication among authors. Once the writing was considered complete, the librarian editor organized and edited the draft in order to create a coherent and consistent paper, then returned this final draft to all authors for their approval. Though coordinating with so many authors is challenging, here we demonstrated that it is possible for a large group of individuals to contribute substantively to an article.

Participants reported that they appreciated having structured roles within the teams. Team leads were also important for the success of the team, though their presence was not necessary for the entire hackathon. For example, inclement weather on the second day prevented one of the team leads from attending, but the team still made progress on pipeline production. Given that members of each team came from diverse backgrounds with experience working with a multitude of different data types and resources, the hackathon promoted innovation through team science and consensus-building. For example, it was essential that each pipeline utilize an appropriate test dataset, but many teams had difficulty with data that were located across multiple repositories or could not be used due to errors in metadata or formatting. Thus, teams had to brainstorm other datasets to use or create new ways to process the data. Because each problem encompassed technical challenges inherent in many biological fields, teams needed to consolidate ideas from each member. This allowed teams to not only transcend the difficult data landscape, but fostered a strong learning environment.

Although the ultimate goal of the hackathon was to solve biological problems, participants emphasized that they appreciated this unique opportunity for career development and networking. Participants with strong backgrounds in computer science effectively mentored those who were less computationally savvy, and those with strong biology backgrounds were able to share insights with those who lacked this expertise. Additionally, the hackathon brought together individuals from different research communities who otherwise may have never met and created the potential for establishing new collaborations. In particular, participants early in their careers were able to meet prominent researchers in various fields and receive helpful training advice from the more senior participants. We anticipate that the participants will share their experiences upon returning to their respective institutes.

The organizers learned some valuable lessons from this event. Surprisingly, although the organizers had kept the goals somewhat loosely structured, participants generally asked for more structure, particularly concerning datasets. In the future, the organizers intend to prepare videos for team members concerning the scientific directions of the projects prior to the event. Other informational materials distributed in advance of the event could help participants learn how to complete tasks that took time away from pipeline development, such as how to locate and download datasets. Specific attention will be paid to using the SRA SDK to process small parts of many genomes simultaneously. One team was unable to complete their pipeline, and other teams were affected by time constraints, so moving some of the preparatory work of locating and downloading datasets would help ensure that the teams had adequate time for more substantive work on the pipelines.

From an institutional perspective, the hackathon was also helpful as a means to test NCBI public data repositories. Over the course of this hackathon, several technical issues with respect to data storage, metadata and corruptions were illuminated. These issues as well as constructive feedback about how NCBI should host data were discussed directly with NCBI Director David Lipman.

Finally, we hope that this hackathon will help to stimulate the community to continue to improve these pipelines. We chose these topics and questions because they are of interest to many biologists and introductory bioinformaticians. Because the data is publicly available, investigators should be able to access the datasets from NCBI in order to replicate the work done in creating these pipelines. We encourage members of the community to extend, expand and alter these pipelines, which are licensed under a Creative Commons Attribution License (CC-BY). We hope that the community will continue working with these pipelines to suit their needs and repost them as they see fit.

## Acknowledgements

Shamira Shallom helped document parts of DNA-seq Team progress. Michelle Dunn, Don Preuss, Julia Oh, and Matt Shirley helped with planning the event. Phi Ngo and other members of the NCBI administrative team provided additional support during the event. Kurt Rodarmer and Wolfgang Goetz provided guidance on the SRA toolkit, and Tom Madden provided guidance about BLAST.

